# Neural patterns of conscious visual awareness in the Riddoch syndrome

**DOI:** 10.1101/2023.04.20.537641

**Authors:** Ahmad Beyh, Samuel E Rasche, Alexander Leff, Dominic ffytche, Semir Zeki

## Abstract

The Riddoch syndrome is one in which patients blinded by lesions to their primary visual cortex can consciously perceive visual motion in their blind field, an ability that correlates with activity in motion area V5. Our assessment of the characteristics of this syndrome in patient ST, using multimodal MRI, showed that: 1. ST’s V5 is intact, receives direct subcortical input, and decodable neural patterns emerge in it only during the conscious perception of visual motion; 2. moving stimuli activate medial visual areas but, unless associated with decodable V5 activity, they remain unperceived; 3. ST’s high confidence ratings when discriminating motion at chance levels, is associated with inferior frontal gyrus activity. Finally, we report that ST’s Riddoch Syndrome results in hallucinatory motion with hippocampal activity as a correlate. Our results shed new light on perceptual experiences associated with this syndrome and on the neural determinants of conscious visual experience.

## INTRODUCTION

The Riddoch syndrome, which constitutes a rich field for studies of visual consciousness, refers to the fact that patients blinded by lesions in their primary visual cortex (area V1) can have a crude but conscious experience of some visual stimuli, prominent amongst which is visual motion. It was first described indifferently by George Riddoch 1 in the aftermath of the Great War but was dismissed by Gordon Holmes ^2^ (for a review of its early history see ^3,4^). The syndrome fell into oblivion until the 1970s when Sanders et al. ^5^ and Weiskrantz et al. ^6^ described a seemingly related syndrome termed blindsight, which was defined as “visual capacity in a field defect in the absence of acknowledged awareness” ^7^, thus contradicting the conscious dimension that Riddoch had written of. It was only in the 1990s that Barbur et al. ^8^ and Zeki and ffytche ^3^ established that hemianopic patients can be conscious of their residual visual capacity.

Imaging studies of such patients ^3,8–10^ led to two conclusions that form the basis of the work reported here. The first is that the conscious experience of visual motion always correlates with activity in V5, an area of the primate visual brain that is specialized for the processing of visual motion ^11,12^; but to be perceived consciously the moving visual stimuli must have certain characteristics, namely be fast moving, be of high contrast, and of low spatial frequency ^3,13^. The second conclusion is that moving visual stimuli that are not perceived consciously also result in weaker but detectable activity within V5.

We hoped that the simple question that we addressed in this work would lead to a simple answer and reveal what it is that dictates a quantitative difference in V5 activity between two states, one in which the patient is conscious of the visual stimulus and its direction of motion, and another in which they are not and can only discriminate motion direction at chance levels ^3^. Two main possibilities, not necessarily exclusive of each other, presented themselves: that the heightened activity is due to the more intense response of cells that are engaged in the conscious perception of visual motion, or that the conscious perceptual state entails the recruitment of an additional population of cells; in the latter instance, the spatial arrangement of activity in V5 should be different in the conscious state. We favoured the latter hypothesis and conjectured that decodable patterns within V5 will only emerge during the conscious experience of visual motion.

We tested this hypothesis in patient ST (not his real initials) who became hemianopic following a posterior cerebral artery stroke that damaged his V1; we assessed him using psychophysics, as well as structural, functional, and diffusion magnetic resonance imaging (MRI). Our results confirmed our main hypothesis – that patterns emerge in V5 only during conscious visual experience – and also led to a more detailed study of a phenomenon which is the opposite of “blindsight”, namely the presence of high perceptual certainty despite chance performance ^3^, leading to the involvement of areas implicated in conflict resolution. Equally important and previously unreported is the appearance of post-stimulatory visual hallucinations in a patient’s blind field, with hippocampal activity as a correlate.

## RESULTS

### Visual assessment of patient ST

ST is a male in his early fifties who experienced a posterior cerebral artery ischaemic stroke resulting in partial loss of vision in his right visual field. Despite this, he showed signs of residual visual processing of motion in his blind field during clinical testing, suggesting that he fits the description of a Riddoch syndrome patient.

Automated, static, binocular Esterman perimetry confirmed that ST was blind in a large portion of his right visual field with sparing of some portions along the meridians (Figure 1). He described the visual disturbance not as a static blackness, but more like a persistent migraine aura that is permanently ‘flickering’ in his lower right visual field, and he insisted that he could distinguish this flicker from motion. He reported seeing shapes and colours in an unclear and fuzzy manner, which he described as a ‘flickering smudge’. We confirmed his report with psychophysical testing in the lab, where he was unable to detect static bright dots of various sizes presented in his perimetrically blind field but saw the same dots if they moved quickly (we tested him at a speed of 10 °/s).

**Figure 1.**
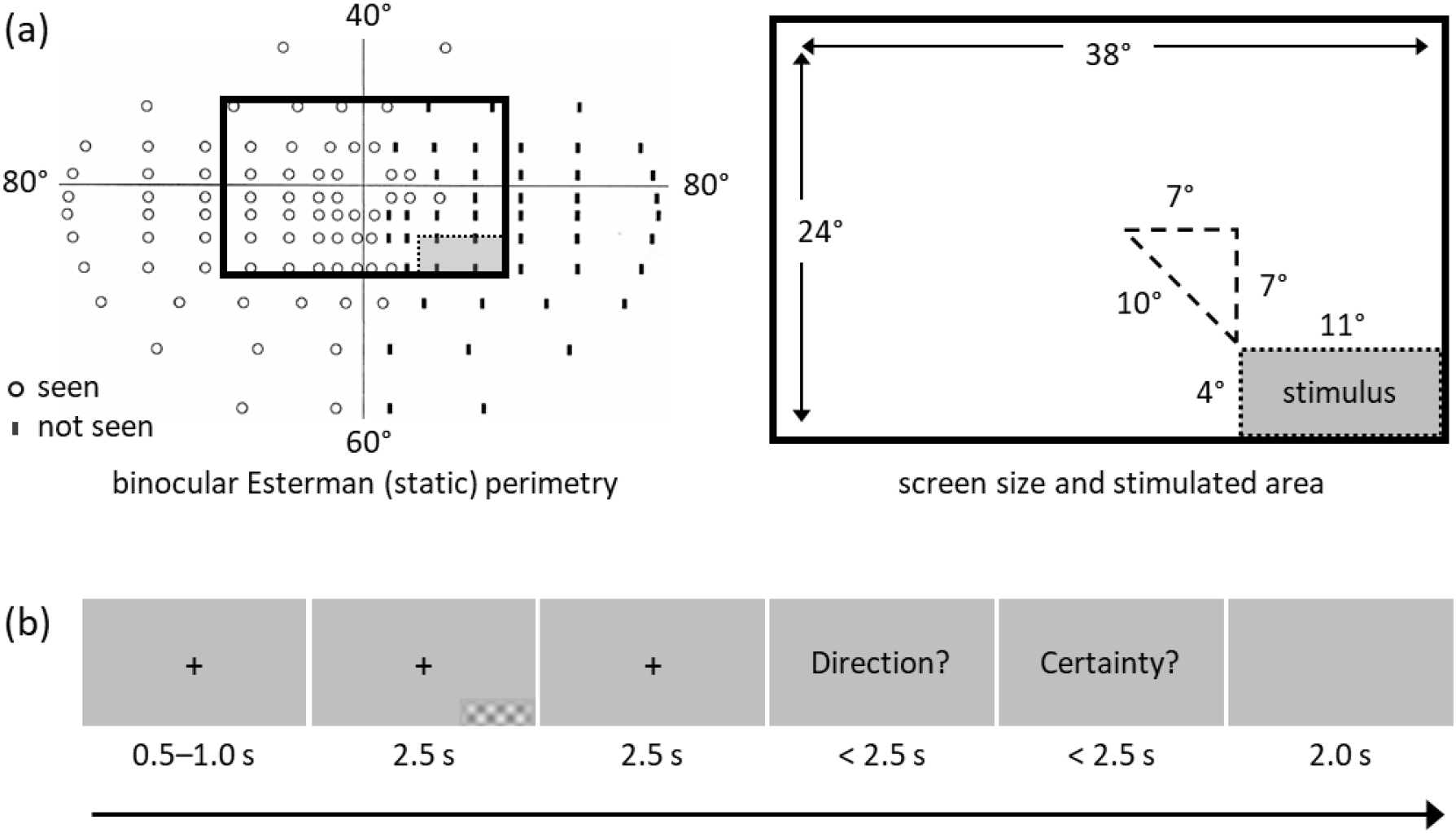
Visual field assessment and visual motion task. **(a)** Visual field assessment delineating ST’s blind field, and the area in which the stimulus was displayed during the fMRI experiment. This part was chosen to ensure that the stimulus did not encroach on the spared parts of the field near the horizontal and vertical meridians. **(b)** The task that ST performed during the psychophysics and imaging experiments. After a short and variable cue, a stimulus appeared for a maximum duration of 2.5 s in ST’s blind field. The stimulus was a drifting achromatic checkerboard which varied in spatial frequency (0.3 or 1.4 cycles/°), contrast (20% or 80%), and speed (1 or 8 °/second), resulting in eight different stimuli. Blank trials on which no stimulus was presented (catch trials) were also included. The patient was then asked to indicate in which direction the checkerboard had moved (upward or downward) in a forced-choice manner, and to indicate his level of certainty about the answer on a three-point scale. The time under each frame in the figure indicates its duration.

### Behavioural results

Our aim was to establish the characteristics of the visual stimulus that ST could perceive consciously. We used achromatic checkerboard stimuli that varied in spatial frequency, contrast, and speed, as well as blank trials (Figure 1). Based on the results of the psychophysics session (Table S1), ST performed the task while undergoing brain imaging.

During the imaging session, ST was very good at discriminating the direction of motion of low spatial frequency drifting checkerboards presented in his blind field (Figure 2 and Table S2). He performed perfectly (100% accuracy) when contrast and speed were high; but he also performed very well with other combinations of contrast and speed as long as the spatial frequency was low (84% accuracy with all low frequency conditions combined). Conversely, his performance was at chance level when presented with high spatial frequency checkerboards, although there were indications, during the psychophysics session, that he could occasionally reach above-chance levels of performance with such stimuli (Table S1). Notably, his certainty level was very high (2.65 ± 0.49 out of 3.00) whenever he was presented with a high speed, high contrast, and high spatial frequency stimulus, despite performing at chance, which made this condition of special interest as it gave him a false sense of confidence. On about half of the blank trials (no stimulus), ST reported moderate to high certainty that he had correctly discriminated the direction of motion of the absent visual stimulus (Figure S1). Based on this result, we divided the blank trials into those with low and those with high certainty, as only the former can be used as a true ‘blank’ condition that does not elicit a conscious percept. Additionally, higher certainty correlated with faster responses as confirmed by a strong negative correlation between certainty and reaction time (*r*_*s*_ = -0.69, *p* < .0001; Figure 2). This supports our observation that ST’s subjective report of certainty was consistent with his experience and could be used as a meaningful metric.

**Figure 2.**
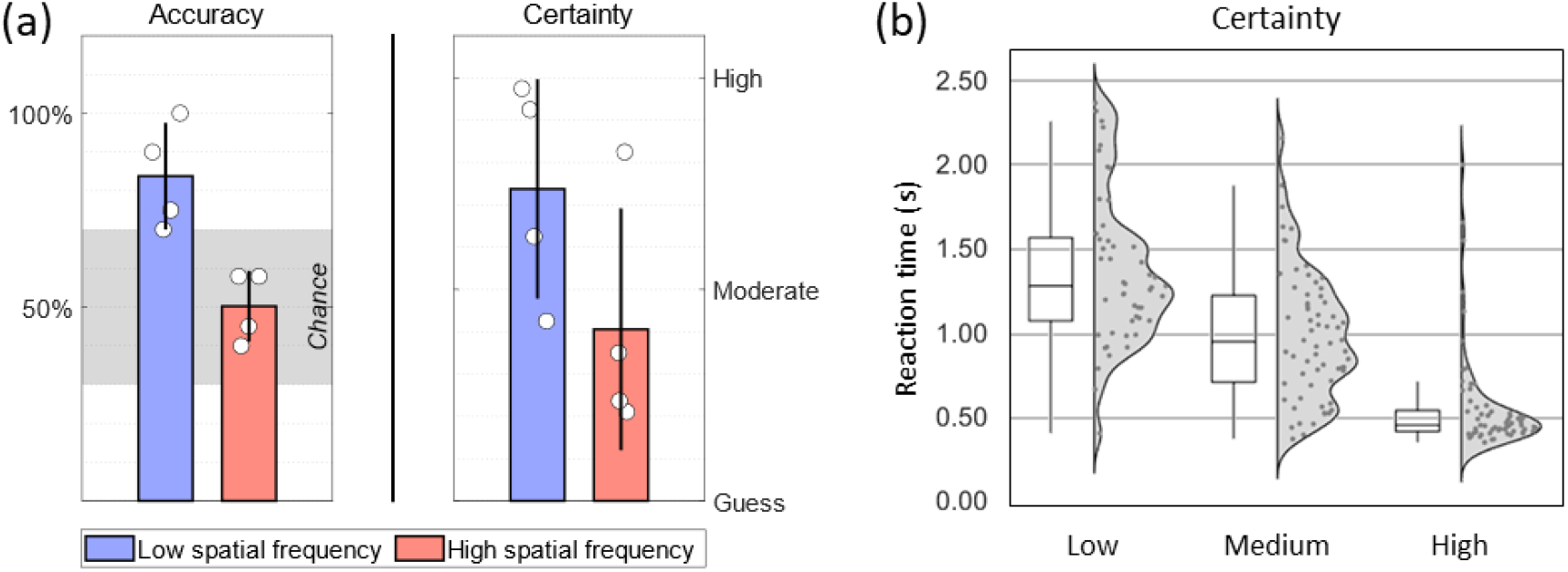
Behavioural results from the visual motion task. **(a)** ST’s performance was highly influenced by the spatial frequency of the checkerboard stimulus: he performed very well with low frequency checkerboards and at chance with high frequency ones. The bars represent the mean accuracy and certainty scores separately for the low and high frequency conditions; the white disks represent the average scores of individual conditions. The shaded grey area in the accuracy plot indicates the 95% confidence interval of chance performance based on the binomial distribution for 20 trials. **(b)** ST’s reaction time to indicate the direction of motion during the fMRI task. A Spearman rank correlation between reaction time and certainty was strong (*r*_*s*_ = -0.69, *p* < .0001), indicating that higher certainty correlated with faster responses.

When ST’s performance is plotted on a psychophysical model adopted from Zeki and ffytche ^3^, it becomes apparent that his perceptual states, induced by the different stimuli, largely fall along the expected continuum between blindness and conscious vision. This further supports the observation that his experience (certainty) and accuracy are closely related (Figure 3).

**Figure 3.**
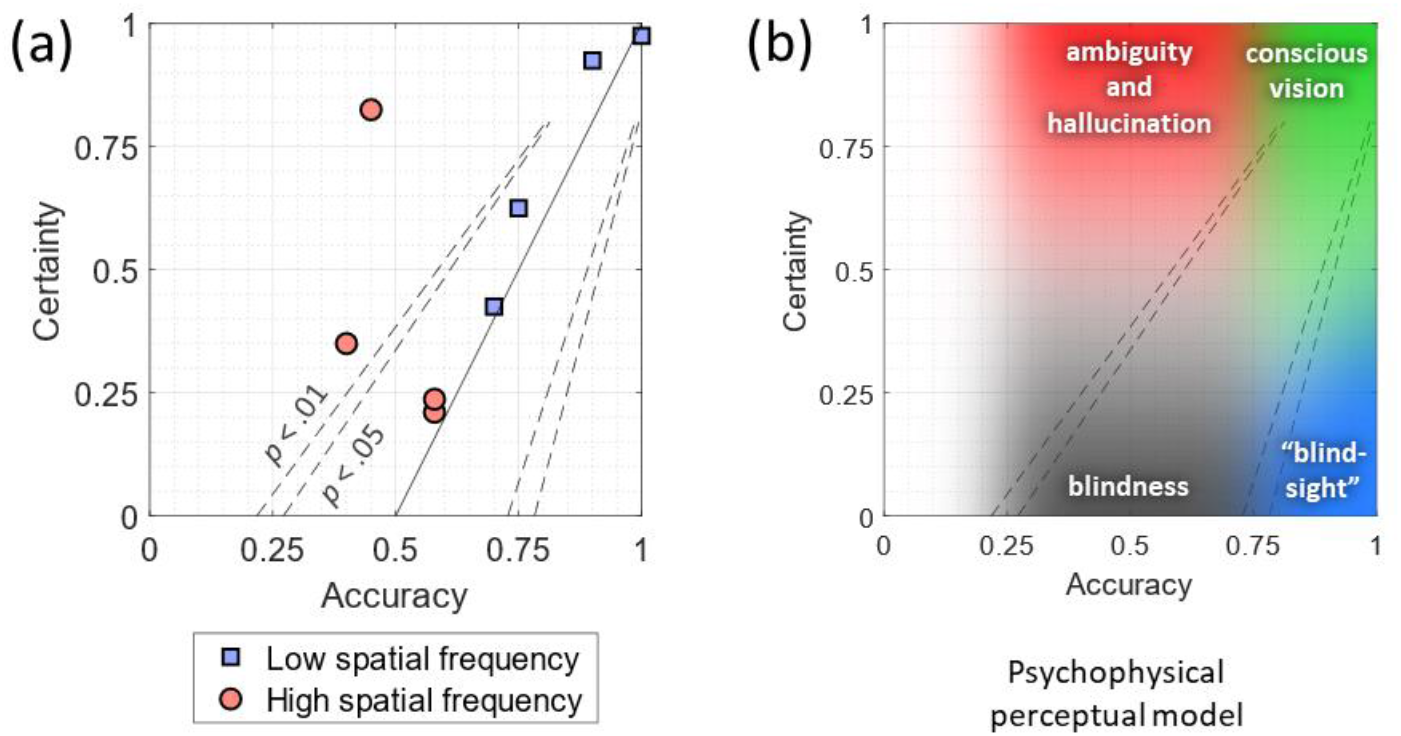
Psychophysical model. **(a)** The solid line represents a psychophysical model that assumes that certainty and accuracy are strongly linked; the dashed lines represent the boundaries of the model under the binomial distribution at *p* < .05 and *p* < .01, calculated for 20 trials per condition. ST’s certainty was congruent with accuracy, except for one strong departure from this trend where ST thought he performed very well but was in fact at chance. **(c)** Various perceptual states placed within the same psychophysical model, showing that ST’s perceptual states largely follow the continuum between blindness and conscious vision, with occasional departures toward ambiguity and hallucination.

### Lesion extent and white matter input to V5

Structural T1w images revealed that ST had a circumscribed lesion (7.27 millilitres in volume) confined to his medial occipital lobe, affecting the calcarine sulcus and pericalcarine grey and white matter (Figure 4). Area V1 was the most affected, but tissue near the occipital pole (subserving central vision) was spared. The surrounding area V2 may also have been affected in part. The lesion extended more into the cuneus than the lingual gyrus and did not approach the locations of areas V3 and V4 ventrally, nor that of V3 dorsally. Importantly, the lesion did not extend laterally enough to reach area V5. A tractography reconstruction of the optic radiations confirmed that ST’s V5 in the lesioned hemisphere remained directly connected with the lateral geniculate nucleus (LGN), and that the microstructure of these connections was indistinguishable from that in the contralesional hemisphere (Figure 4).

**Figure 4.**
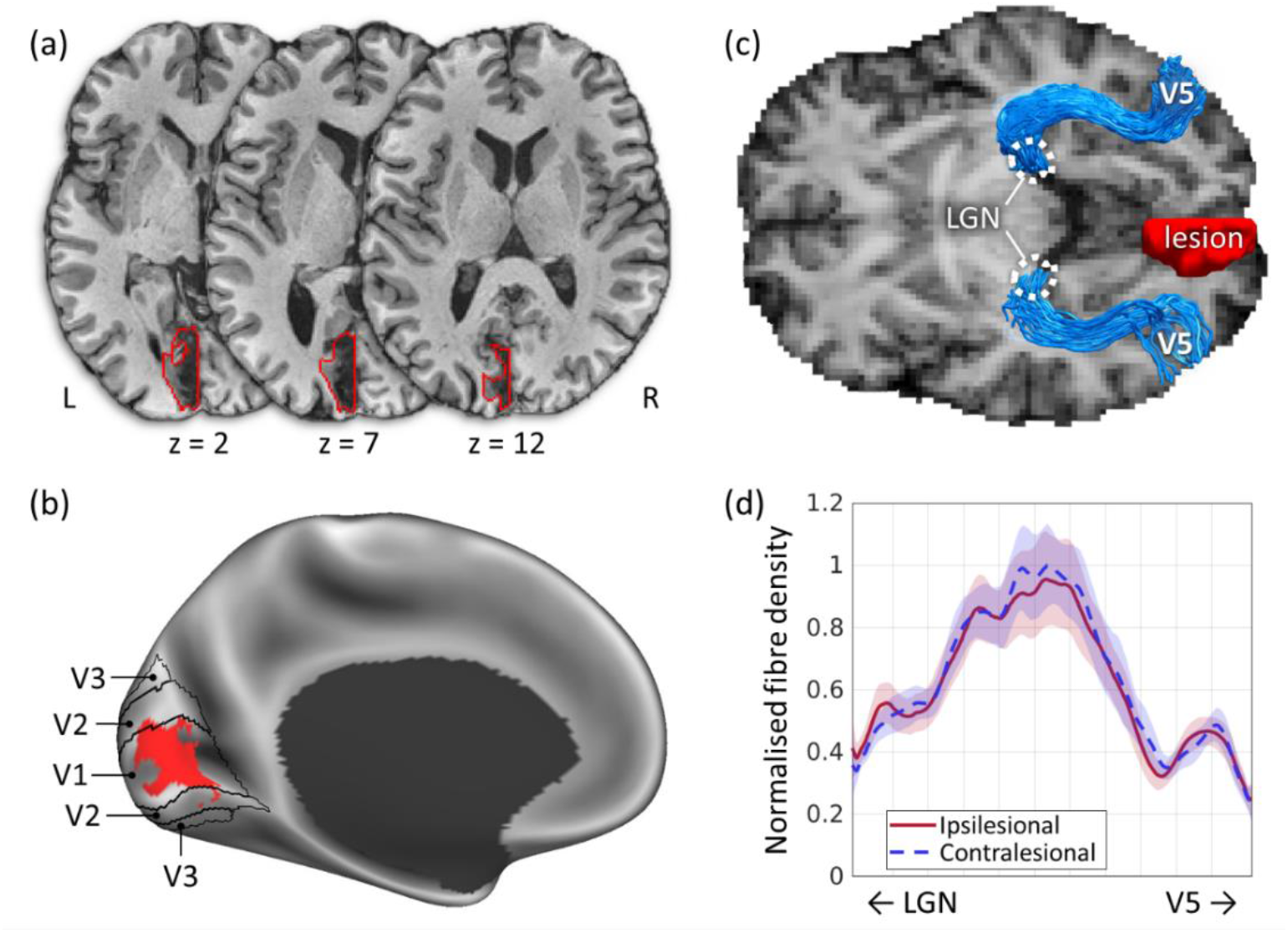
ST’s lesion extent and reconstructed optic radiations terminating in V5. **(a)** Axial slices through ST’s T1w structural image showing that the lesion (red contour) was confined to V1 and did not extend laterally to affect V5. **(b)** The lesion displayed on a canonical brain surface, showing the full extent of its cortical reach. Notice the crescent shape within the posterior calcarine sulcus that corresponds to the spared visual field along the horizontal meridian in the perimetry plot (Figure 1). **(c)** Tractography reconstruction of the optic radiations connecting the lateral geniculate nucleus (LGN) with area V5, in both hemispheres, displayed against the anisotropic power map derived from the diffusion data. The location of V5 was determined from the fMRI task. **(d)** Microstructural comparison of the LGN-V5 connections in the ipsilesional and contralesional hemispheres based on the hindrance modulated orientational anisotropy (HMOA), a proxy for fibre density. No difference is observed between the two hemispheres.

### Functional imaging results

We conducted various univariate and multivariate analyses to assess brain activity related to different perceptual states while ST performed the visual motion fMRI task; we focus here on the four conditions of main interest (others are reported in Figure S2).

The first condition (*Seen*) represents conscious vision (high accuracy and high certainty); here, the stimulus had a low spatial frequency, high contrast, and high speed. The second condition (*Not seen*) represents blindness, i.e., the inability to consciously perceive a stimulus presented in the blind field (chance discrimination and low certainty); the stimulus in this condition was of high spatial frequency, low contrast, and low speed. The third condition (*Ambiguous*) represents a false sense of confidence which is incongruent with performance (high certainty despite chance discrimination); in this case the stimulus had a high spatial frequency, high contrast, and high speed. Finally, the fourth condition (*Hallucinated*) represents imagined vision, i.e., the experience of seeing a moving stimulus despite none being presented; this condition consisted of blank trials for which ST reported moderate to high certainty in discriminating “motion” direction.

The results of these combined analyses revealed several clusters in the occipital, inferior frontal, orbitofrontal, and cingulate cortices, as well as the tail of the hippocampus. A summary of the main results is presented in Figure 5 and detailed results are reported in Figure S2 and Table S3.

**Figure 5.**
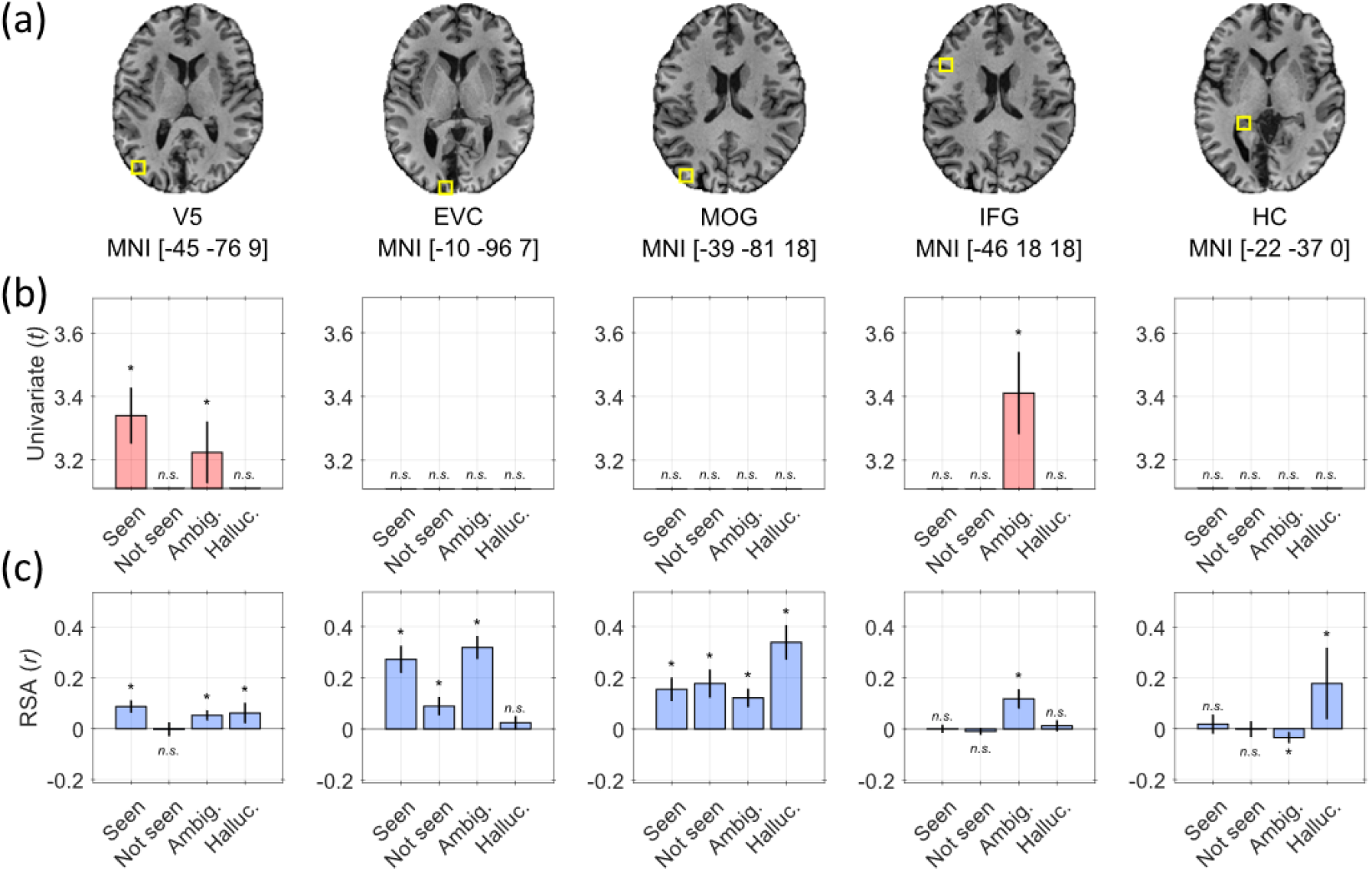
Univariate and multivariate activity during the visual motion task. Functional MRI during the visual motion task revealed various regions implicated in processing conscious or unconscious visual motion perception. **(a)** The location (yellow square) and MNI coordinates of each region of interest (ROI) are shown on an anatomical T1w image of ST’s brain. **(b)** Each panel shows the t-statistic of the change in activity in the corresponding ROI in (a) relative to the ‘low certainty blank’ condition from a univariate analysis. **(c)** Each panel shows the strength of multivariate activity patterns in the same ROIs and for the same comparisons as in (b). The regions are V5, early visual cortex (EVC), middle occipital gyrus (MOG), inferior frontal gyrus (IFG), and the tail of the hippocampus (HC).

#### Univariate analyses

The univariate analysis, which compared activity associated with each condition to activity associated with low certainty blanks, revealed increased activity in V5 during the *Seen* condition, and in both V5 and the inferior frontal gyrus (IFG) during the *Ambiguous* condition.

#### Multivariate analyses

For the representational similarity analysis (RSA) ^14^, our model assumed that neural patterns would be similar for trials of the same condition (e.g., *Seen*), and different from those of the low certainty blank trials; it further assumed that the latter would not share a common pattern (Figure S3). According to this model, patterns emerged in V5 only when ST reported a high level of certainty, i.e., during the *Seen, Ambiguous*, and *Hallucinated* conditions, but not when he was unconscious of the visual stimuli and failed to discriminate motion direction (*Not seen*) (Figure 5).

Although our main aim was to investigate activity patterns in V5, RSA revealed the involvement of several other regions in the various conditions (Figure 5). We observed patterns in early visual cortex (EVC, areas V2/V3) when a stimulus was present, regardless of ST’s level of certainty. Patterns emerged in (a) the middle occipital gyrus (MOG) for all four conditions but were most similar during the *Hallucinated* condition; (b) in the inferior frontal gyrus (IFG) only during the *Ambiguous* condition, which is in line with univariate activity in that region; (c) and in the tail of the hippocampus (HC), which showed strong pattern similarity only during the *Hallucinated* condition (we ensured that this is not mislabelled LGN activity, see Figure S4). This suggests that activity in these regions was too weak to be visible in the univariate analysis, but that they were nonetheless directly involved.

## DISCUSSION

We began this research with a simple question and ended up, unexpectedly, on shores we had not intended to visit. We enquired into patient ST who fits the profile of a classical Riddoch syndrome patient in that, despite becoming hemianopic after a lesion to V1, he retained the ability to perceive visual motion consciously in his blind field. We assessed him using psychophysics and MRI, but obtained a complex set of results that speaks not only to the Riddoch syndrome, but also to the neural correlates of various experiential states, including conscious visual perception, ambiguity, and hallucination, as defined above.

### A lesion in V1 sparing subcortical input to V5

Structural imaging revealed that ST’s lesion is confined to V1 in the left hemisphere, and tractography confirmed that V5 in the same hemisphere receives direct input from the LGN. These findings are in accordance with the expected anatomy of a Riddoch syndrome patient based on previous reports ^3,15^.

### Behavioural determination of conscious awareness of visual motion in the blind field

We specifically tailored the stimuli to create conditions in which ST could discriminate visual motion direction easily and consciously, conditions at the threshold of conscious awareness, and ones in which he is blind to visual motion. Our behavioural results showed that we successfully induced this spectrum of experiential states in ST; our results are in agreement with those obtained by Zeki and ffytche ^3^ in patient GY, with perceptual states largely falling along the continuum between blindness and conscious vision, and with a tight link between performance and awareness (Figure 3). Thus, we find no evidence in ST of the dissociation between performance and awareness after V1 damage as described by “blindsight”, where a patient can unconsciously discriminate visual motion direction with high accuracy ^16,17^. In fact, several reports have since shown that such findings can be fully explained by a methodological bias in how patients are asked about their experience, and that discrimination and conscious awareness are very closely related ^3,18–21^.

The variety of stimuli that we used did, however, induce experiential states associated with the Riddoch syndrome that have not been explored extensively, including ambiguous and hallucinatory states (Figure 3). For example, when the stimulus was of high spatial frequency, high speed, and high contrast, ST reported a high level of certainty despite chance discrimination; and during some blank trials where no stimulus was presented, he reported being moderately to highly certain of discriminating the direction of motion of the non-existent visual stimuli.

### Consciously seen motion correlates with decodable neural patterns in V5

When we used stimuli of low spatial frequency, high speed, and high contrast – stimuli associated with good discrimination and conscious awareness – the results of the univariate analysis were straightforward; they showed that there was heightened activity in V5. The multivariate analysis revealed distinct neural patterns for these stimuli in V5. In addition, though absent in the univariate analysis, presumably because of relatively weak activity, significant pattern similarity was also found in EVC and the MOG. The patterns in EVC fell at the border between areas V2 and V3; although neither has been shown to be specifically or exclusively involved in visual motion, V2 has been called a “distributor” area because, among its compartments of specialised cells, is one where directionally selective cells are concentrated (the thick cytochrome oxidase stripes) which project anatomically to V5 ^22,23^. Further, V3 is largely dominated by an M input, responds to visual motion, and has good concentrations of directionally selective cells in it, ranging from 12% ^24^ to 40% ^25^ in the macaque, and has been observed in the human brain to respond to both first and second-order motion ^26^. The MOG patterns, on the other hand, can be attributed to its role in categorising visual inputs in general, as has been demonstrated by previous reports ^27,28^.

### V5 activity is an essential complement to consciously seen motion

When the stimulus was of high spatial frequency, low speed, and low contrast, it was unperceivable to ST; he was unable to discriminate its motion direction and reported low certainty. These stimuli engage the magnocellular pathway weakly and the parvocellular one strongly, and thus depend on a healthy V1 or early visual cortex for their perception, rather than on V5; it is therefore not surprising that ST, with his lesion in V1, was unable to perceive these stimuli. Consistent with this, there were no significant univariate activations anywhere in the brain compared to low certainty blanks. However, we did find significant pattern similarity for this type of stimulus in the MOG and EVC. While the role of the MOG may be seen as an attempt to categorise even the weakest visual input, spared EVC, which still receives direct subcortical input, can exhibit very weak but decodable activity that is not sufficient to evoke a conscious percept ^29^. Therefore, moving stimuli may give rise to neural activity in medial visual areas, but unless this is associated with V5 activity, they remain unseen by Riddoch syndrome patients.

Furthermore, previous reports on patients ^3^ and controls ^30^ have demonstrated that visual motion that is not perceived consciously can activate V5, though to a lesser extent than motion that is seen consciously. Our results do not contradict this; in fact, we observe that some moving stimuli that are not perceived consciously and are associated with chance performance can occasionally activate V5, but they are not associated with patterns in it (Figure S2). Therefore, we only observe patterns in V5 in association with consciously perceived motion, even if univariate activity can be detected in it.

### High certainty with chance discrimination correlates with activity in the inferior frontal gyrus

A stimulus of high spatial frequency, high speed, and high contrast evoked a false sense of certainty in ST; that is, he reported being highly confident in perceiving and correctly discriminating the direction of motion despite his ability to do so remaining at chance. We refer to this condition as ambiguous, because there was a certainty which only he perceived ^31^; it is a phenomenon strongly reminiscent of *gnosanopsia* ^3^. Note that ST’s performance was at chance level, indicating that he was not consistent in his imperception; he was not, e.g., consistently reporting the opposite of the correct motion direction, as has been reported in a case of akinetopsia ^32^, nor was he consistently responding with a single motion direction (e.g., upward on all trials). It would therefore be more accurate to label this as a directionally “bistable” percept, which is well supported by the neuroimaging results. In fact, there was increased activity, as well as decodable patterns, in the IFG, which has been shown to play a crucial role in resolving perceptual conflict and stabilizing visual awareness when different interpretations are equally valid ^33–35^. Additional areas also showed distinct neural patterns for this type of stimulus, namely V5, EVC, and the MOG. This indicates that these visual stimuli were indeed perceived as moving, but that their direction of motion was ambiguous, requiring input from the IFG to reach a resolution.

### Hallucinatory motion in the Riddoch syndrome correlates with hippocampal activity

Another interesting finding is that, during blank trials where no stimulus was presented, ST occasionally reported having moderate-to-high certainty of having seen a moving stimulus and discriminating its direction of motion. We consider these occurrences to be hallucinations or imagery of visual motion, though they may not necessarily be clear or vivid; therefore, they may be regarded as minor rather than complex hallucinations ^36^. This is supported by the imaging results, which showed univariate activity in bilateral orbitofrontal cortex (Table S3), and multivariate patterns in V5, the MOG, and the tail of the hippocampus. The fact that there were patterns in V5 is a strong indication that these trials were associated with a visual motion percept as previous work has shown that the visual category of a hallucination correlates with activity in the visual areas specialised for that type of content ^37^. Additionally, the MOG and hippocampus have been implicated in imagery and the retrieval of visual perceptual content from recent memory ^38,39^. Although we did not explicitly address this question, we suspect that the hallucinations observed here are task-induced, in that they may be brought upon by strong expectations about encountering visual motion immediately after the cue at the start of each trial ^40^. In fact, several recent reports have proposed that hippocampal neural representations generate cued predictions about upcoming sensory events, modulating activity in sensory cortex in a predominantly top-down fashion ^41–44^.

### Concluding remarks

Only experiential states in which ST reported some degree of awareness of motion direction showed distinct neural patterns in V5. One possible criticism may be that the neural activity associated with each experiential state is driven by low-level stimulus properties and is difficult to disentangle from activity associated with conscious experience. However, the fact that neural patterns emerged in V5 only during conditions with conscious experience, be it driven by a clear, ambiguous, or hallucinated percept, speaks to a common thread connecting these conditions *despite* the differences in low-level features.

On the other hand, the various experiential states engaged different sets of areas along with V5, some of which are more generally involved in conscious perceptual processing and not necessarily restricted to visual motion. The activity in these areas, which was relatively weak and therefore not easily demonstrable by univariate analysis, could be decoded through to multivariate analysis. This highlights the importance of tackling patient cases and group studies with multiple analytical tools, as this may reveal mechanisms of conscious perception which are otherwise occult. Our results also highlight the complex continuum of perceptual experiences in patients ‘blinded’ by cortical lesions, and their ability to tell us about the neural mechanisms of conscious perception, within and beyond visual cortex.

In summary, our results confirmed the supposition of decodable patterns emerging in V5 during the residual conscious vision of subjects ‘blinded’ by lesions in V1. But surprisingly, they also opened new and unexpected fields of enquiry – into the relationship between certainty and conscious experience, between blindness and visual hallucinations, and ultimately into the meaning of these decodable patterns that correlate with conscious experience.

## ACKNOWLEDGEMENTS

This work was supported by a Leverhulme Trust grant to Semir Zeki (RPG-2020-022).

## AUTHOR CONTRIBUTIONS

Conceptualization, A.B., S.E.R., S.Z.

Methodology, A.B., S.E.R., D.f., S.Z.

Software, A.B., S.E.R.

Formal Analysis, A.B., S.E.R.

Investigation, A.B., S.E.R., A.F., D.f., S.Z.

Resources, A.F., S.Z.

Writing – Original Draft, A.B., S.E.R., S.Z.

Writing – Review & Editing, A.B., S.E.R., A.F., D.f., S.Z.

Visualization, A.B., S.E.R.

Funding Acquisition, S.Z.

## METHODS

### Resource availability

#### Lead contact

Further information and requests for resources should be directed to the Lead Contact, Ahmad Beyh.

#### Materials availability

This study did not generate new unique reagents.

#### Data and code availability

Custom MATLAB analysis code available from the corresponding author upon request.

### Method details

#### Patient

ST was referred to our study via the National Hospital for Neurology and Neurosurgery in London. He gave informed written consent to participate in our study, which had been approved by the Yorkshire & The Humber - South Yorkshire Research Ethics Committee (NHS Health Research Authority) and UCLH/UCL Joint Research Office (protocol number 137605).

#### Visual motion task

Our aim was to establish the characteristics of the visual stimulus that ST could perceive consciously. We began by using achromatic checkerboard stimuli which varied in spatial frequency (0.3 or 1.4 cycles/°), contrast (20% or 80%), and speed (1 or 8 °/second), resulting in eight different stimuli. Blank trials during which no stimulus was presented (catch trials) were also included to assess ST’s baseline response to the task. The checkerboard was confined to ST’s blind visual field and moved either upward or downward on each trial (Figure 1). The edges of the stimulus were blurred to avoid a sharp boundary between it and the surrounding grey background, and its mean luminance was matched to that of the background. The task was programmed in PsychToolbox 3 ^45^, running in Matlab (MathWorks, Natick, MA).

Figure 1 shows a schematic of the task. Each trial started with a cue (fixation cross) which lasted for 0.5-1.0 s. Next, the stimulus appeared and lasted for a total duration of 2.5 s, including 0.5 s of fade-in and 0.5 s of fade-out, followed by a rest period of 2.5 s. Then, two questions were presented sequentially: the first asked ST to indicate the direction of motion following a two-alternative forced-choice design (2AFC); the second asked ST about his level of certainty on a three-level scale. The maximum time allowed to respond to each question was 2.5 s, but the task moved on as soon as a response was recorded. Finally, a blank screen (grey background without a fixation cross) was displayed for 2.0 s as a rest period between trials.

#### Procedure for psychophysical testing

ST viewed the stimuli at a 60 cm distance while seated and resting his chin on a support. First, we confirmed the extent of the blind field as revealed by perimetry with the use of a laser pointer. ST was asked to indicate whether he saw a red dot appear on the screen while fixating on a cross at the centre of the screen. Next, he was presented with an achromatic checkerboard of varying contrast, spatial frequency, and moving at varying speeds in in his blind field, to examine the threshold of his residual visual capacities. The stimulus was restricted to the lower right portion of his visual field (7° below the horizontal meridian and 7° to the right of the vertical meridian) and subtended 12° in width and 5° in height; it faded in and out, with each fade lasting for 0.5 s, and was shown at full opacity for 1.5 s, amounting to a 2.5 s total stimulation time. Blank trials were also included. The stimulus presentation was followed by a 2.5 s blank screen, after which the questions were presented. Using a keyboard, ST was asked to indicate the direction of motion in a two alternative force choice (2AFC, up or down) paradigm and his certainty (awareness) on a three-point scale, 1 corresponding to ‘complete guess’, 2 to ‘somewhat certain of the direction’, and 3 to ‘very certain of the direction’. The questions were presented for 3 s each but were removed from the screen as soon as a response was registered. This again was followed by a blank screen, lasting for 2 s. Three runs of 30 trials were performed.

In the second session, ST was again asked to perform the same task with the same checkerboard confined to the same portion of his blind field. This time we conducted a 2×2×2 design, with either low or high speed (1 or 8 °/s), contrast (20% or 80%) and spatial frequency (0.3 or 1.4 cycles/°). Each run contained four trials of each combination and six blanks. We conducted six runs of 38 trials, amounting to a total of 24 trials per condition and 36 blanks (228 total trials).

Performance on each condition was determined by taking the fraction of correct responses in direction discrimination. The statistical significance of his performance was determined using a binomial test. The average certainty score was also calculated for each condition.

#### Procedure for testing during the imaging session

Based on the results of the initial psychophysics session (Table S1), ST performed the task while undergoing brain imaging. The task was divided into four runs; during each, the eight stimuli were presented randomly, five times each, in addition to five blank trials. This amounted to 45 trials per run, or 180 trials in total, with each stimulus (including blanks) being presented 20 times. During each trial, ST indicated the direction of motion with his right hand and his certainty with his left hand using a customised button-box.

#### Imaging acquisition

We acquired structural, functional, and diffusion MRI data on a 3T Siemens Prisma scanner (Siemens Healthcare GmbH, Erlangen, Germany) with a 64-channel head coil. The structural images were based on a 3D magnetisation-prepared accelerated gradient echo (MPRAGE) sequence: repetition time (TR) = 2.53 ms; echo time (TE) = 3.34 ms; flip angle = 7°; matrix of 256×256; field of view = 256 mm; voxel size = 1×1×1 mm^3^.

fMRI data were based on the blood oxygen level dependent (BOLD) signal, measured with a 2D T2*-weighted Echo Planar Imaging (EPI) sequence: volume TR = 3360 ms; TE = 30 ms; flip angle = 90°; ascending acquisition; matrix of 64×64; voxel size = 3×3×3 mm^3^; 48 slices. A total of four fMRI runs were acquired. Field mapping images were also acquired using a dual-echo gradient echo sequence to assist with susceptibility distortion correction.

Diffusion MRI data were based on a 2D spin-echo EPI sequence: TR = 3500 ms; TE = 61 ms; flip angle = 88°; matrix of 110×110; voxel size = 2×2×2 mm^3^; 72 slices; multiband factor of 2; in-plane acceleration factor of 2. Images were acquired with three diffusion shells: 30 diffusion directions at b = 500 s·mm^-2^, 60 directions at b = 1500 s·mm^2^, and 90 directions at b = 2500 s·mm^-2^. Additionally, 16 b = 0 s·mm^-2^ were interleaved throughout the acquisition, and seven b = 0 s·mm^2^ volumes were acquired with the reverse phase encoding polarity to correct for susceptibility distortions.

#### T1w MRI pre-processing

The T1w image was skull-stripped using *optiBET* ^46^, bias field corrected using the *N4* tool ^47^, and rigidly aligned, using *flirt* ^48^, to the 1 mm MNI T1w brain template ^49^ as a substitute for AC-PC alignment. This aligned image served as the anatomical reference for subsequent pre-processing and analysis steps. Additionally, the T1w image was normalised to the MNI template through affine and diffeomorphic non-linear transformations (SyN algorithm) computed with ANTs ^50^. A manually delineated lesion mask was used to exclude the lesioned tissue during the normalisation step.

#### fMRI pre-processing

The first four volumes of each fMRI run were discarded to allow the scanner to reach steady state. The remaining images were corrected for motion and slice-timing differences using *SPM12* (http://www.fil.ion.ucl.ac.uk/spm/software/). The corrected images were then simultaneously corrected for geometric distortions (based on the acquired field map) and aligned to the T1w image using FSL’s *epireg* tool ^48,51^, while maintaining the voxel size at 3×3×3 mm^3^. This produced the final fMRI time series images which were used in subsequent analyses.

#### Diffusion MRI pre-processing

Raw DWI data were first corrected for noise and Gibbs ringing artefacts ^52,53^. A magnetic susceptibility field was then calculated using *topup* ^54^ based on b = 0 s·mm^-2^ images acquired with opposite phase-encoding. All images were subsequently corrected for motion and eddy current distortions using *eddy* ^55^ with outlier (signal dropout) slice replacement ^56^, incorporating the *topup* field into this step. The anisotropic power map was derived from the pre-processed data using StarTrack (www.mr-startrack.com) and used to calculate a rigid affine transformation (six degrees of freedom) to the T1w image with *flirt*. The rigid transformation was then applied to the diffusion data (kept at a 2 mm voxel size) with a spline interpolation to produce the final set of pre-processed images. The diffusion gradients were also rotated at this stage using the same transformation matrix.

#### Univariate fMRI analysis

Various categorical comparisons were made to assess the activity related to different perceptual and certainty states. Each combination of factors of the 2×2×2 design was considered as a separate condition, which were compared to one another and to the blank condition. As ST reported some level of certainty on nearly half the blank trials, indicating that he had seen something moving, we decided to divide the blank trials into two distinct conditions: ‘low certainty blanks’ included trials receiving a rating of 1 (i.e., total guess), and ‘high certainty blanks’ included trials receiving a rating of 2 or 3 (i.e., somewhat to very certain).

For the univariate analysis, the BOLD time series images were spatially smoothed with a Gaussian kernel of a full width at half maximum (FWHM) of 4.5 mm. The time series were entered into a general linear model (GLM) in *SPM12* with a single task effect (stimulus presentation) and six motion correction parameters as nuisance regressors. A contrast image was generated to compare each condition to the ‘low certainty blank’ condition, and this contrast was entered into a t-test to assess its statistical significance at each voxel. All resulting statistical images were thresholded at a voxelwise significance level of *p* < .001.

#### Representational similarity analysis

For the multivariate analysis, the BOLD time series were first entered into a GLM in *SPM12*, without any spatial smoothing, with a single task effect (stimulus presentation) and six motion correction parameters as nuisance regressors. Each trial was modeled as a separate condition, thereby generating a parameter estimate (beta image) for each.

To investigate whether certainty in perceiving the motion direction of a stimulus in the blind field was accompanied by specific spatial patterns of neural activity, we used representational similarity analysis (RSA) ^14^. The 20 beta maps of each condition and the nine beta maps of the low certainty blank trials were selected and a whole-brain searchlight analysis was performed using cubic regions of interest (ROI) of 3×3×3 voxels. For each searchlight ROI, the Pearson correlation distance, *d*, was calculated between the neural patterns associated with these trials, for each pair of trials, as follows:

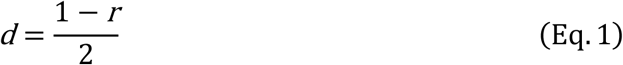

where *r* is the Pearson correlation coefficient; the division by two was performed to rescale *d* to the range [0-1]. This was done to simplify the interpretation and visualisation of the metric: *d* = 0.0 corresponds to a full correlation between the neural patterns of two trials (i.e., *r* = 1.0); *d* = 0.5 corresponds to the absence of any correlation (i.e., *r* = 0.0); and *d* = 1.0 corresponds to the two trials having completely anti-correlated patterns (i.e., *r* = -1.0).

Once the Pearson distances were calculated for each pair of trials, neural representational dissimilarity matrices (RDMs) were generated to capture the (dis)similarity between pairs of trials in each searchlight ROI. The Pearson correlation distance is mainly sensitive to the spatial pattern of brain activity and is insensitive to the overall BOLD signal amplitude change in a given ROI ^14^. Given that the aim here is to find a specific pattern of activity regardless of amplitude, the Pearson correlation distance is the preferred distance metric for our purposes (unlike, e.g., the Euclidean distance which would also record overall magnitude changes like the univariate framework) ^14^.

To test whether the similarity was significant only for the trials of a given condition (and not for those of the low certainty blanks), the RDM was calculated for each searchlight ROI and compared to a model RDM (Figure S3). The correlation between the neural and model RDMs was assessed with the Spearman rank correlation, only using the elements in the lower triangle of the RDMs (excluding the diagonal).

For a given condition, the model RDM that we tested (Figure S3) assumed a high similarity in the activity patterns associated with the trials of that condition (i.e., *d* = 0.0), and no similarity for the trials of the low certainty blanks, or between the patterns of that condition and those of the low certainty blanks (i.e., *d* = 0.5). No pairs of trials were expected to have anti-correlated patterns (i.e., *d* = 1.0) as this would be a strong assumption to make.

#### Tractography

The diffusion data were used to reconstruct the optic radiations connecting ST’s lateral geniculate nucleus (LGN) to area V5 of his visual cortex. The data were modelled with spherical deconvolution based on the damped Richardson-Lucy algorithm ^57,58^ in StarTrack, according to the following parameters: fibre response *α* = 1.5; number of iterations = 350; amplitude threshold *η* = 0.0015; geometric regularisation *ν* = 16.

A probabilistic dispersion tractography approach was followed to explore the full profile of the fibre orientation distribution function (fODF) in each voxel. This approach follows the principal fibre orientations indicated by the fODF local maxima, as well as the directions captured by other vertices of the fODF that convey information about various local fibre orientations ^58^. Fibre tracking was performed according to the following parameters: minimum HMOA threshold = 0.0025; number of seeds per voxel = 2000; maximum angle threshold = 40°; minimum fibre length = 50 mm; maximum fibre length = 150 mm. This was done using a seed region of interest in the LGN obtained from a published atlas ^59^. The resulting tractogram was imported into TrackVis (http://trackvis.org/) where manual cleaning was performed and streamlines terminating in V5 were selected.

For the along-tract microstructural analysis, each streamline was divided into 100 equal segments between its LGN and V5 terminations, and the mean HMOA value of all streamlines was calculated for each segment along with the 95% confidence interval.

#### Quantification and statistical analysis

The statistical tests used were described in detail in the Methods Details subsection for each assay and in the relevant figure legends.

