## Supplemental Material for "Neural patterns of conscious visual awareness in the Riddoch syndrome"

Ahmad Beyh [1,4,*] & Samuel E Rasche [1,4], Alexander Leff [2], Dominic ffytche [3], Semir Zeki [1,*]

[1] *Laboratory of Neurobiology, University College London, London, UK*
[2] *UCL Queen Square Institute of Neurology, University College London, London, UK*
[3] *Department of Old Age Psychiatry, Institute of Psychiatry, Psychology and Neuroscience, King's College London, London, UK*
[4] *These authors contributed equally*

This document contains supplemental material related to the main manuscript and contains:

- Results from the psychophysics testing session.
- Additional behavioural results from the fMRI task.
- Additional fMRI results.
- Table of the fMRI analysis results.

**Table S1. Behavioural results from the psychophysics session.**

| | Low spatial frequency (LF) | | High spatial frequency (HF) | |
|---|---|---|---|---|
| | **Low contrast (LC)** | **High contrast (HC)** | **Low contrast (LC)** | **High contrast (HC)** |
| **Low speed (LS)** | A: 75%*, C: 2.58 | A: 92%*, C:2.83 | A: 50%, C: 1.96 | A: 83%*, C: 2.50 |
| **High speed (HS)** | A: 92%*, C: 2.88 | A: 96%*, C: 2.83 | A: 79%*, C: 2.04 | A: 63%, C: 2.67 |

Accuracy and certainty scores in visual motion direction discrimination for stimuli varying in contrast, speed, and frequency. *A* represents accuracy in percentages, *C* represents the average certainty rating on a scale from 1-3. For the blank condition, certainty was 1.6 on average. Each condition was presented 24 times, except for the blank condition, which was presented 36 times. This amounted to a total of 228 trials. *Significantly different from chance performance ($p < .05$) determined from the binomial distribution for 24 trials.

**Table S2. Behavioural results from the fMRI session.**

| | Low spatial frequency (LF) | | High spatial frequency (HF) | |
|---|---|---|---|---|
| | **Low contrast (LC)** | **High contrast (HC)** | **Low contrast (LC)** | **High contrast (HC)** |
| **Low speed (LS)** | A: 70%*, C: 1.85 | A: 75%*, C:2.25 | A: 55%, C: 1.42 | A: 40%, C: 1.70 |
| **High speed (HS)** | A: 90%*, C: 2.85 | A: 100%*, C: 2.95 | A: 55%, C: 1.47 | A: 45%, C: 2.65 |

Accuracy and certainty scores in motion direction discrimination for stimuli varying in contrast, speed and frequency. *A* represents accuracy in percentages, *C* represents the average certainty rating on a scale from 1-3. *Significantly different from chance performance ($p < .05$) determined from the binomial distribution for 20 trials.

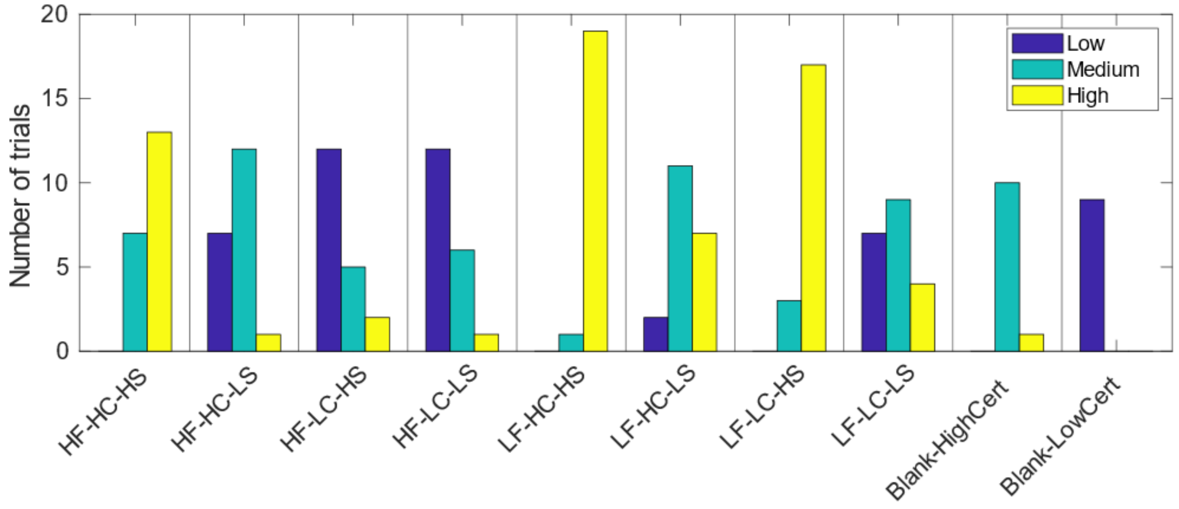


**Figure S1. Certainty ratings across conditions.**
The plots show the distribution of ST's certainty ratings for each task condition. The names of the conditions along the x-axis describe the stimulus properties. For example, LF-HC-HS corresponds the stimulus with low spatial frequency (LF), high luminance contrast (HC), and high speed (HS). Blanks are shown here as two conditions to highlight the difference between high and low certainty blank trials, which was the basis of how we defined the *Hallucinated* condition and the true *Blank* condition.

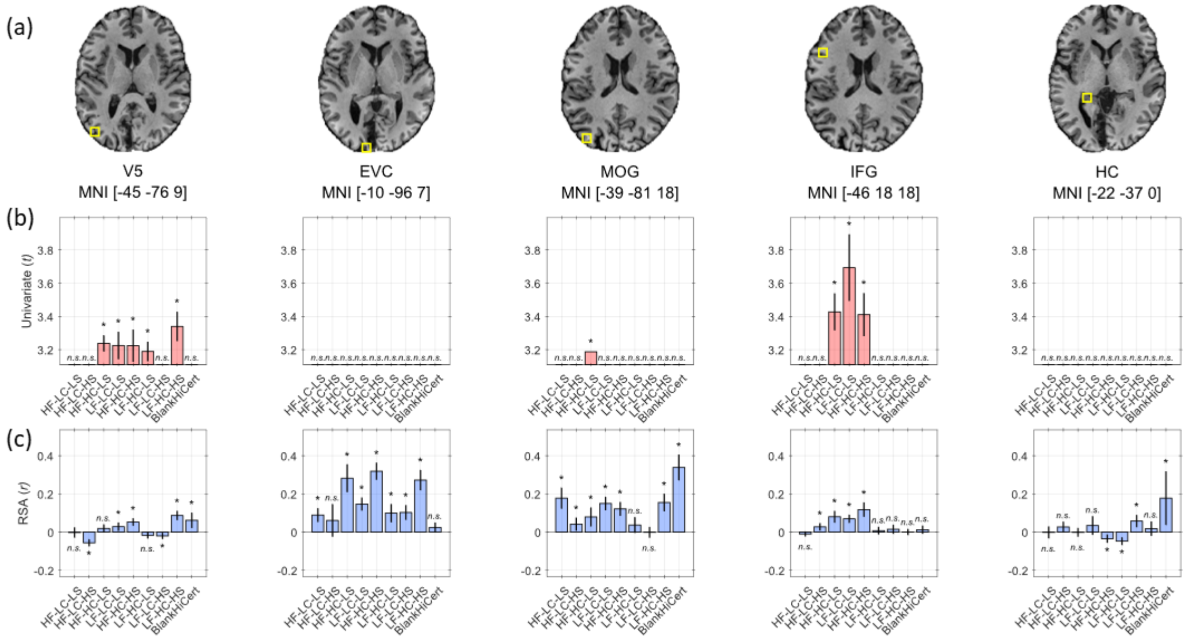


**Figure S2. Univariate and multivariate results for all conditions of the visual motion task.**
**(a)** The location (yellow square) and MNI coordinates of each region of interest (ROI) are shown on an anatomical T1w image of ST's brain. **(b)** Each panel shows the t-statistic of the change in activity in the corresponding ROI in (a) relative to the 'low certainty blank' condition from a univariate analysis. **(c)** Each panel shows the strength of multivariate activity patterns in the same ROIs and for the same comparisons as in (b). The regions are V5, early visual cortex (EVC), middle occipital gyrus (MOG), inferior frontal gyrus (IFG), and the tail of the hippocampus (HC). Naming convention for the conditions as in Figure S1.

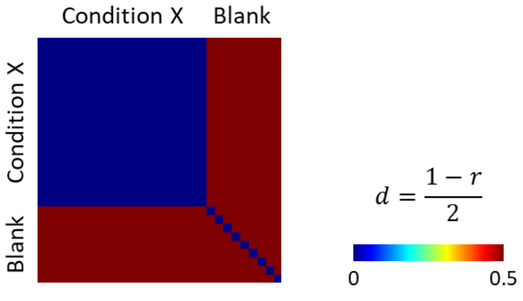


**Figure S3. Model RDM for RSA.**
The model representational dissimilarity matrix (RDM) used for the representational similarity analysis (RSA). The model, which is based on the Pearson correlation distance shown on the right, assumes a similar spatial activity pattern for trials of a given condition, and no similarity between them and those of the low certainty blank condition, or among the latter.

**Table S3. Results of the univariate fMRI analysis.**

| Cluster and/or region | # voxels | *p* | *t* | Coordinates (mm) | | |
|---|---|---|---|---|---|---|
| | | | | x | y | z |
| ***Low spatial frequency conditions > low certainty blanks*** | | | | | | |
| L V5 | 31 | 0.000 | 3.48 | -48 | -78 | 9 |
| ***LF-HC-HS > low certainty blanks*** | | | | | | |
| L V5 | 38 | 0.000 | 3.67 | -45 | -78 | 9 |
| L precentral gyrus | 35 | 0.000 | 3.53 | -36 | -15 | 60 |
| ***High spatial frequency conditions (not including HF-HC-HS) > low certainty blanks*** | | | | | | |
| No significant activations. | | | | | | |
| ***HF-LC-LS > low certainty blanks*** | | | | | | |
| No significant activations. | | | | | | |
| ***HF-HC-HS > low certainty blanks*** | | | | | | |
| L inferior frontal gyrus (IFG) | 66 | 0.000 | 3.64 | -45 | 27 | 18 |
| L IFG | | 0.000 | 3.63 | -45 | 15 | 24 |
| R cingulate gyrus mid-body | 11 | 0.000 | 3.38 | 12 | 15 | 36 |
| R precentral gyrus | 9 | 0.001 | 3.31 | 45 | 6 | 33 |
| V5 | 14 | 0.001 | 3.27 | -42 | -75 | 12 |
| ***High certainty blanks > low certainty blanks*** | | | | | | |
| R orbitofrontal cortex | 8 | 0.000 | 3.41 | 21 | 21 | -15 |
| L orbitofrontal cortex | 9 | 0.000 | 3.36 | -21 | 24 | -15 |
| R cingulate gyrus mid-body | 10 | 0.001 | 3.28 | 12 | 18 | 39 |

For each condition of interest, the clusters of significant activity are reported with their corresponding peak t-statistic and MNI coordinates. All results are thresholded at $p < 0.001$, and only clusters of five voxels or more are reported.

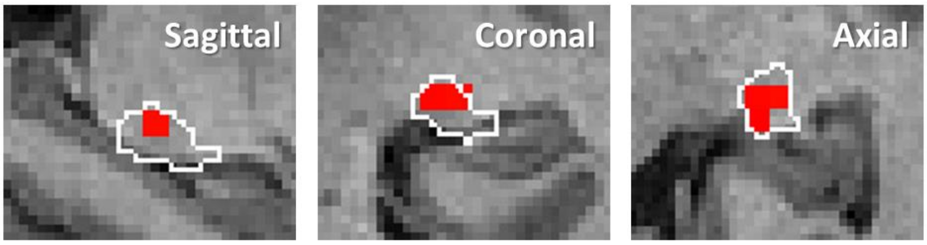


**Figure S4. Location of the lateral geniculate nucleus.**
The white contour shows the atlas-based anatomical location of the LGN in the left hemisphere, and the red voxels correspond to the functionally defined LGN obtained from an F contrast using all the visual task conditions, during which a stimulus was displayed in the right 'blind' visual field. In line with this, no voxels were activated in the right (contralesional) hemisphere.
